# Characterizing the genetic polymorphisms in 370 challenging medically relevant genes using long-read sequencing data from 41 human individuals among 19 global populations

**DOI:** 10.1101/2022.08.03.502734

**Authors:** Yanfeng Ji, Jiao Gong, Fritz J Sedlazeck, Shaohua Fan

## Abstract

Numerous challenging medically relevant genes (CMRGs) cannot be adequately investigated using next-generation sequencing, hindering the detection of functional variation among these genes. In this study, long-read sequencing data from 41 human individuals across 19 populations were analyzed using the current version of the human reference genome assembly (GRCh38) and a telomere-to-telomere assembly of the human genome (T2T-CHM13). After excluding 142 CMRGs containing windows with a depth of coverage (DoC) significantly deviating from the average DoC value of proteincoding regions in the GRCh38 (138) or T2T-CHM13 (47) assemblies, 179 and 263 CMRGs exhibited copy number variation (CNV) signal in GRCh38 and T2T-CHM13, respectively. In addition, 451 high-impact short variants were detected in 188 CMRGs. Further, some genetic alterations were individual- or continental-superpopulation-specific, suggesting a strong need to consider genetic background differences in future genetic testing and drug design studies. Finally, side-by-side comparisons of short variant calls in CMRGs using NGS and LRS data from 13 samples indicated that 15.79% to 33.96% of high-impact short variants in different individuals could only be detected using LRS data. The results described herein will be an important reference for future clinical and pharmacogenetic studies to further improve precision medicine.

## INTRODUCTION

Next-generation sequencing (NGS) has become a routine diagnostic analytical tool in many clinical laboratories due to its low cost and high efficiency for large-scale analysis. Indeed, increasing numbers of clinical laboratories are transitioning from gene panels that target a limited number of known loci to whole-exome sequencing (WES) that has been used for clinical testing and pharmacogenetic studies (1–3). For example, a WES investigation of 208 pharmacogenes in 60,706 unrelated individuals observed abundant functional rare variants in pharmacogenes (4).

Although NGS, especially WES, can be widely applied in clinical studies, it also has some strong limitations (5, 6). First, due to its poor performance (7, 8), NGS (both WES and WGS) has not been used in routine genetic testing of copy number variation (CNV), which has a clear role in drug-related genes by altering drug metabolism, transportation, and response (9). Further, detection of short variant, including single nucleotide variations (SNVs) and short insertions and deletions (InDels), in segmental duplications and highly repetitive regions is also challenging for NGS-based methods, leading to some WES studies completely avoiding these regions (10). This is particularly problematic for clinical studies since numerous highly challenging medically relevant genes (CMRGs) are located in repetitive or highly polymorphic regions of the human genome (11–13). For example, previous studies have shown that 17,561 pathogenic variants of CMRGs are difficult to investigate using NGS-based methods (11). Therefore, challenges remain in characterizing genetic variation in CMRGs using NGS data.

Long-read sequencing (LRS) technologies including Pacific Biosciences (Pacbio) and Oxford Nanopore Technology (ONT) platforms have greatly expanded our understanding of human genomes (14–19). LRS technologies allow sequencing up to > 10 kbp read length and the ability to sequence through repetitive regions. Consequently, LRS has been widely used to study complex variation in human genomes (14–20). Recent studies have shown that ~68% of structural variants (SVs) that were detected with LRS cannot be detected using NGS (14) and LRS data yield better short variant (including SNV and InDel) calling accuracy, especially for variants located in difficult-to-map genomic regions based on NGS datasets (21). LRS data has also been used to generate a telomere-to-telomere assembly of the human genome (T2T-CHM13) that has significantly improved the assembly of the remaining sequences in centromeric, telomeric, and segmentally duplicated regions of the current human reference genome version GRCh38 (13, 22–24).

LRS genomic data from a Jewish trio were used by the Genome in a Bottle (GIAB) group to benchmark 273 CMRGs in the human reference genomes GRCh37 and GRCh38 (25). These analyses identified assembly errors in six total MRGs of GRCh37 (*MRC1* and *CNR2*) and GRCh38 (*CBS, CRYAA, KCNE1* and *H19*) (25). Further, a significant increase in variant calling accuracy of NGS data was observed after excluding six false duplicate CMRGs (25). This pattern was also confirmed by the T2T consortium (13). Further, the T2T-CHM13 assembly facilitated the detection of CMRG structural variants (13). Nevertheless, a comprehensive investigation of CMRG genetic variation across global populations using LRS remains lacking.

In this study, the assembly status of 370 CMRGs on both the T2T-CHM13 and GRCh38 assemblies was evaluated by leveraging LRS data across 19 human populations comprising 41 samples with African, American, South Asian, East Asian, and European ancestries. Genetic polymorphisms were then investigated within these CMRGs including SNVs, InDels, and CNVs. These analyses strengthen our understanding of the genetic diversity of CMRGs, while also providing a critical reference for future clinical and pharmacogenetic studies (26, 27). The results presented herein also highlight the need for more advanced methods to characterize and identify these complex genes.

## MATERIALS AND METHODS

### CMRGs in public databases

A dataset comprising 370 CMRGs generated by the GIAB Consortium was analyzed here (25). Briefly, CMRGs were obtained using annotations from the Online Mendelian Inheritance in Man (OMIM) (28), Human Gene Mutation Database (HGMD)(29), ClinVar (30), and GeneCard (www.genecards.org) databases.

### Human reference genomes

The human reference genome GRCh38 (version hg38_GCA_000001405.15) (31) was retrieved from the UCSC genome browser website (32), while T2T-CHM13 version 1.1 (22) was retrieved from https://s3-us-west-2.amazonaws.com/human-pangenomics/T2T/CHM13/assemblies/chm13.draft_v1.1.fasta.gz. Only autosomes were included in the analysis.

### Annotation of gene and segmental duplication regions in GRCh38 and T2T-CHM13

The gene annotations of GRCh38 were analyzed from the database record https://ftp.ebi.ac.uk/pub/databases/gencode/Gencode_human/release_38/gencode.v38. ann otation.gff3.gz, in addition to the T2T-CHM13v1.1 record from https://s3-us-west-2. amazonaws. com/human-pangenomics/T2T/CHM13/assemblies/annotation/chm13.draft_v1.1.gene_annotation.v4. gff3.gz.

### LRS data from global populations

LRS data from 37 samples analyzed in previous studies (14, 15, 33) were downloaded from public data repositories (Supplementary Table 1). The LRS data for CHM13 was also retrieved from the NCBI SRA database (accession number: SRX789768).

### LRS mapping and depth of coverage (DoC) calculations

The LRS data were mapped to the GRCh38 and T2T-CHM13 assemblies using Winnomap (34) version 2.03 with default parameters. DoC was measured using the mosdepth program (version 0.3.1) with parameters -b 100 -Q 20, by calculating DoC per 100 bp using unambiguously mapped reads (35).

To account for DoC differences across windows per individual and across samples, normalization was conducted such that the DoC of each window was divided by the genome-wide average DoC value per sample.

### Evaluating the assembly of 370 CMRGs in the GRCh38 and T2T-CHM13 assemblies

The assembly qualities of 370 CMRGs in the T2T-CHM13 and GRCh38 assemblies were evaluated based on coverage distributions. Assuming a random distribution of sequencing reads across genomes (22), regions that significantly deviated from the average DoC of 17,337 protein-coding genes (either DoC < average-3*SD or DoC > average+3*SD) were identified after normalization of the T2T-CHM13 and GRCh38 assemblies, respectively. Regions with significantly low DoC (DoC < average-3*SD) are likely to be regions where long-read sequencing data cannot be unambiguously mapped. These regions may include long repeat regions or those highly similar to other regions (*e.g*., segmental duplications and satellite repeats), in addition to misassembled regions and false duplications within the assembly (25).

In contrast, regions with significantly high DoC (> average + 3*SD) are likely to be regions that collapsed multiple haplotypes into a single consensus sequence during assembly. Previous studies have observed that collapsed regions increase false-positive variant calling since reads from multiple locations are mapped to a single region (36).

### Detection of CNV regions and short variants in global populations

CMRGs were excluded that contained windows with DoC values significantly deviating from the average DoC value for protein-coding genes in the GRCh38 or T2T-CHM13 assemblies.

Potential CNV regions in each sample were identified using the same strategy described above for detecting regions with abnormal DoC in the T2T-CHM13 and GRCh38 assemblies. CNV regions observed in most of the samples (non-reference allele frequency > 95%) were considered likely donors of the reference genome carrying a minor allele (15) in global populations. The CNV variability of a CMRG was calculated as the percentage of windows exhibiting a CNV signal across 41 samples.

The mapping results of the global samples were used as input for short variant (single nucleotide variation and short insertion and deletion) calling. To minimize the impact of excessive sequencing coverage on short variant calling accuracy (37), ten samples with >60x coverage (HG02011, HG02818, HG03065, HG03371, NA19983, NA12329, HG02492, HG03009, HG03683, and HG04217) (Supplementary Table 1) were down-sampled to ~60x coverage prior to short variant calling using the Samtools program (version 1.12) (38) with the -s parameter. PEPPER (21) (version 0.7) was used to detect short variants for samples sequenced with the Pacbio HiFi or ONT platforms. For samples sequenced using Pacbio CLR technology, Clair (version 2) (39) was used for the variant calling model trained for CLR data. The “PASS” variants from the PEPPER or Clair programs were used in further analyses.

Functional annotations of short variants of the CMRGs were identified using the Ensembl Variant Effect Predictor (VEP) program in the offline mode (40).

Due to an average of two individuals per population, we characterized the CNV polymorphism at individual-, intra-continental-, and inter-continental-superpopulation levels.

## Results

Long read sequencing (LRS) data for 41 samples across five ethnicities (including African (14), East Asian (8), South Asian (6), European (5), and American (8) ancestries) as well as CHM13 were sequenced with >30x coverage using multiple sequencing technologies including Pacbio CLR, Pacbio HiFi, and ONT (ultra-long and 1D sequencing) platforms (Supplementary Table 1). Comparison of depth of coverage (DoC) values per 100 bp windows based on the LRS mapping data for CHM13 were first used to evaluate the concordance of the T2T-CHM13 and GRCh38 assemblies. Within the GRCh38 assembly, 2,043,520 windows significantly deviated from the average DoC of the protein-coding gene regions compared to the T2T-CHM13 assembly (435,196 windows) (Supplementary Tables 2-3), consistent with previous studies documenting a significant improvement of the T2T-CHM13 assembly over the GRCh38 assembly (22).

Windows with abnormal DoC values in the CMRGs across the T2T-CHM13 and GRCh38 assemblies were also categorized (Figure 1A). One (one window) and 11 (601 windows) CMRGs contained windows with significantly elevated DoC values across the protein-coding gene regions that were specific to the T2T-CHM13 and GRCh38 assemblies (Supplementary Table 4), respectively. In addition, nine CMRGs contained windows with DoC values significantly greater than the average DoC values of proteincoding gene regions in both the T2T-CHM13 and GRCh38 assemblies (Supplementary Table 4). For example, the analyses suggested that a ~1,700 bp region (Chr5:1,201,154-1, 202,853 in T2T-CHM13; Chr5:1,293,347-1,295,046 in GRCh38) overlapped the first two exons of Telomerase Reverse Transcriptase (*TERT*) and is likely to be collapsed in the T2T-CHM13 and GRCh38 assemblies (Figures 1B-C). *TERT* encodes a catalytic subunit of the telomerase enzyme that maintains telomere structures by adding small, repeated segments of DNA (TTAGGG) to the ends of chromosomes during cell division. Previous studies have shown that *TERT* mutations are associated with genetic diseases like dyskeratosis congenita autosomal dominant 2 that is characterized by progressive bone marrow failure, in addition to the clinical combination of reticulated hyperpigmentation of the upper chest and/or neck, dysplastic nails, and mucosal leukoplakia (41, 42).

**Figure 1.**
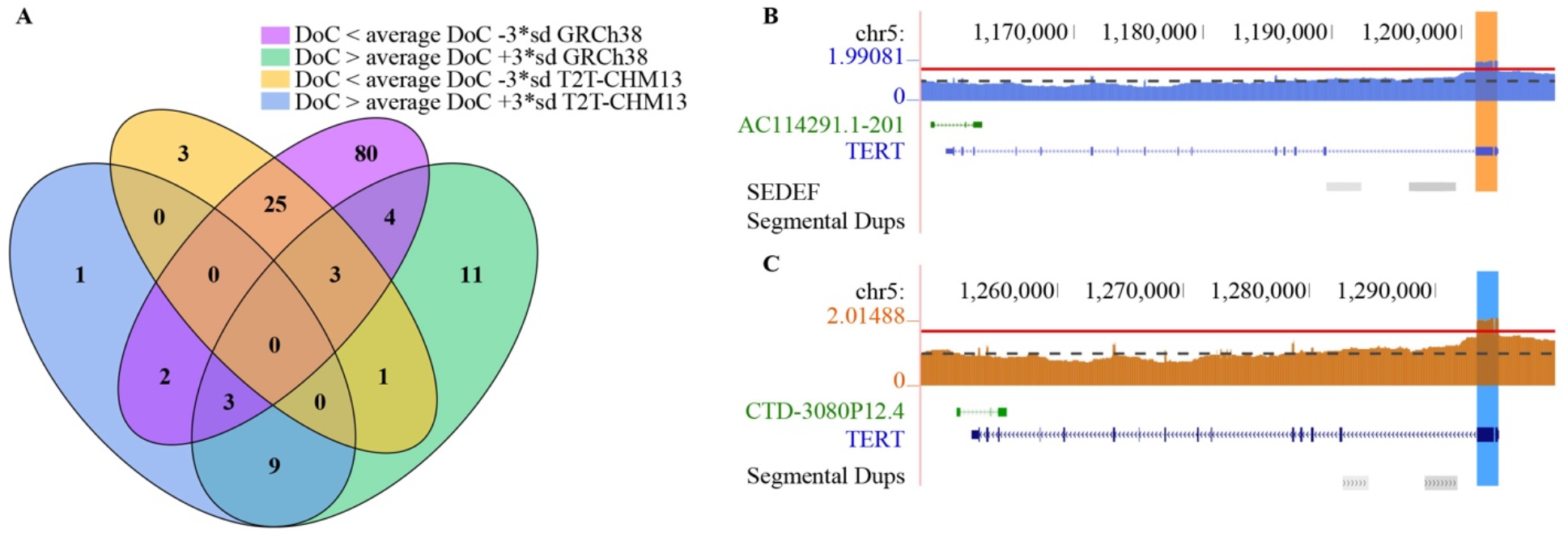
Assessing CMRGs in the GRCh38 and T2T-CHM13 assemblies using PacBio HiFi data from the CHM13 assembly. A: Numbers of CMRGs that contain windows with depth of coverage (DoC) values significantly deviating from the average DoC values of protein-coding gene regions in the GRCh38 and T2T-CHM13 assemblies. B-C: Windows with significantly greater DoC values than the average DoC values of the protein-coding gene regions that overlap with the first two exons of *TERT* in both the GRCh38 (B) and T2T-CHM13 (C) assemblies.

Furthermore, three (79 windows), 80 (3,143 windows), and 25 (1,639 windows in T2T-CHM13 and 1,310 windows in GRCh38) CMRGs were identified that contain windows with DoC significantly lower than the average DoC of protein-coding gene regions and were specific to T2T-CHM13, GRCh38, or were shared by both assemblies (Figure 1A; Supplementary Table 4). A total of 3,693 windows within 56 CMRGs exhibited significantly lower coverage than the average in the GRCh38, while no window in the T2T-CHM13 assembly exhibited such a pattern. This result is potentially due to assembly problems for GRCh38, given that >96% of the windows with zero DoC were identified in or within 100 kbp of the segmental duplication regions (39.24%, 1,449 windows) and the remaining assembly gaps (57.03%, 2,106 windows) in the GRCh38.

Interestingly, the kringle IV type 2 (KIV-2) regions of the *LPA* locus, a regulator of plasma Lp(a) levels, exhibited significantly greater DoC values than the average DoC of the protein-coding gene regions of GRCh38, but significantly lower DoC than the average for protein-coding gene regions of T2T-CHM13. The KIV-2 repeat is ~5.5 kbp long and varies from 5-50+ copies among individuals. Six and 23 copies of the KIV-2 repeat are present in the GRCh38 (43, 44) and T2T-CHM13 (24) assemblies. Consequently, the assessment of this crucial, complex gene is considerably more difficult for the GRCh38 assembly. A 15 kbp region (chr6:161875872-161890671) of the KIV-2 repeat region of T2T-CHM13 exhibited a DoC that was significantly lower than the average DoC of the protein-coding gene regions when realigning its own Pacbio data, suggesting the Pacbio HiFi sequence data may not be able to fully resolve such a long repeat region.

*SHANK2* and *SEC63* contain windows with DoC values significantly greater than the average DoC for protein-coding gene regions in the T2T-CHM13 assembly and windows with zero DoC values in the GRCh38 assembly (Supplementary Figure 1, Supplementary Table 4). Note that the windows in different assemblies do not correspond to each other (Supplementary Figure 1). The regions with zero DoC in *SHANK2* resulted from a remaining assembly gap, but two zero DoC regions in the *SEC63* overlapped with one SINE and one SVA transposable element that are likely to be specific to the GRCh38 donor.

Finally, the analyses indicated that four CRMGs including *DUX4, SMOC2, SNTG2*, and *STK11* contained windows with DoC significantly greater and lower than the average DoC of the protein-coding gene regions in the GRCh38 assembly (Supplementary Table 4).

Overall, both assemblies likely contain windows with DoC that significantly deviate from the average DoC of protein-coding gene regions, but a significant improvement in the assembly of CMRG was observed for the T2T-CHM13 assembly when compared to the GRCh38 assembly. A total of 232 and 323 CMRGs that contain windows with abnormal DoC were used for the GRCh38 and T2T-CHM13 assemblies, respectively, for further analyses of SNVs, InDels, and CNV polymorphisms across 41 global population samples.

### CNV polymorphisms in CMRGs of the global populations

Based on the DoC analyses, 16,424 windows were identified in 179 CMRGs of the GRCh38 assembly (77%) that exhibited CNVs signals, in addition to 31,243 windows in the 263 CMRGs in the T2T-CHM13 assembly (81%) (Supplementary Tables 5-6). When excluding CNV windows in the segmental duplication regions, 10,124 windows in 154 CMRGs (66%) and 20,728 windows in 225 CMRGs (70%) exhibited CNV signals in the GRCh38 and T2T-CHM13 assemblies, respectively (Figure 2A, Supplementary Tables 5-6). Notably, 94.81% of the CMRGs (146 out of 154) with CNVs in the GRCh38 assembly were also identified in the T2T-CHM13 assembly (Figure 2A). Further, the CNV variability, which was defined as the percentage of windows showing CNV signals in a CMRG, of 146 CMRGs in both GRCh38 and T2T-CHM13 (Figure 2B) were observed near the diagonal, suggesting excellent congruence of the CNV analyses based on different assemblies. Finally, *LCE3B* exhibited the greatest variability among 146 CMRGs that exhibited CNV signals in both the GRCh38 and T2T-CHM13 assemblies (Figure 2B). Deletion of *LCE3B* is strongly associated with psoriasis (45) and exists at a high frequency in Eurasians, but not in Africans (46).

**Figure 2.**
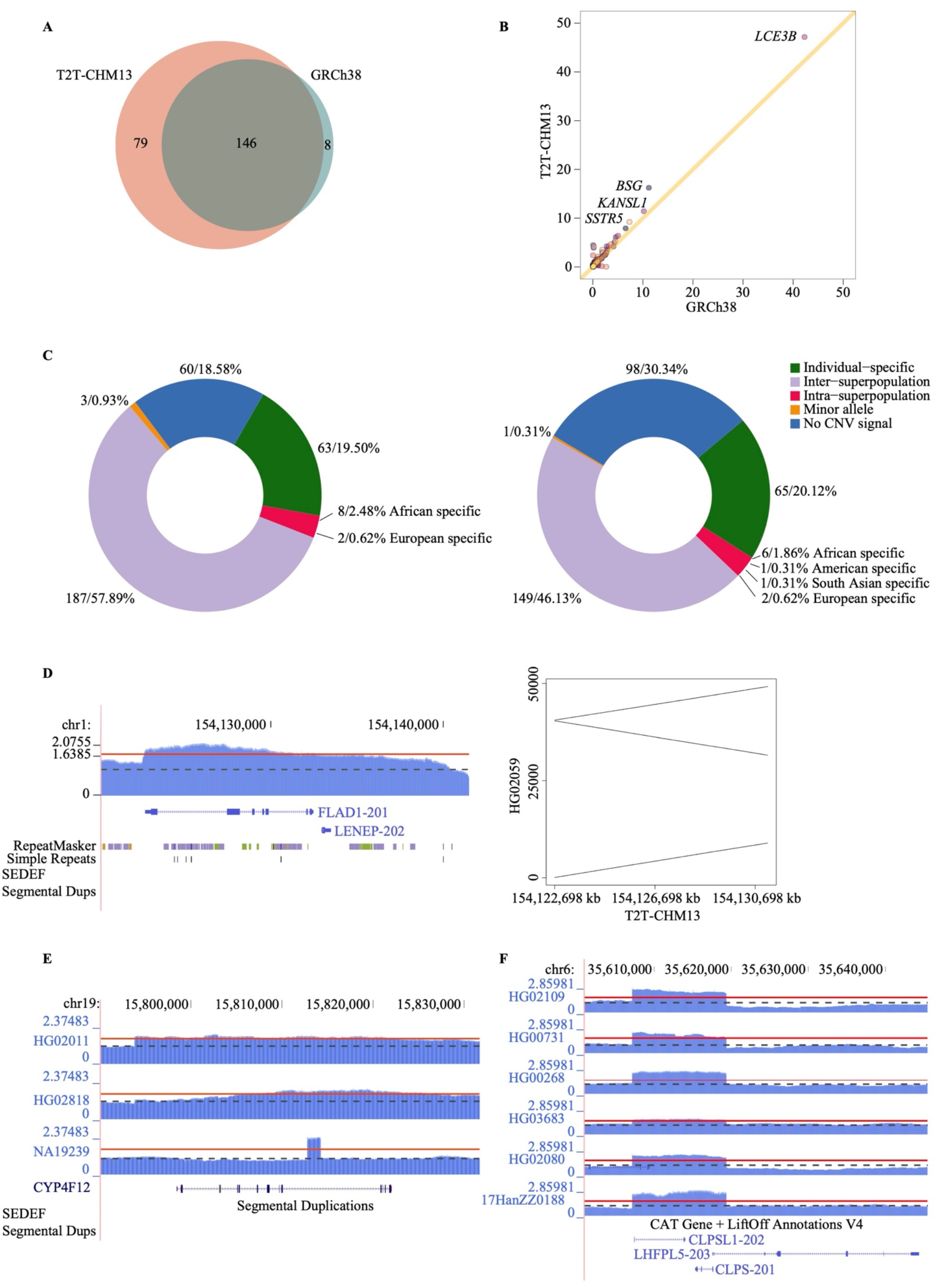
Summary of CMRGs with CNV signals in the global populations. A: Overlap of 233 CRMGs exhibiting CNV signals among global populations based on the GRCh38 (154 CRMGs) and T2T-CHM13 (225) assemblies. In addition, 94.81% of the CMRGs with CNV signals in the GRCh38 assembly were also identified in the T2T-CHM13 assembly. B. CNV variability of 146 CMRGs across 41 samples. CMRG variability was defined as the percentage of windows in a CMRG exhibiting CNV signals. The analyses were based on 146 CMRGs with CNV signals in both the GRCh38 and T2T-CHM13 assemblies. The X- and Y-axes indicate the percentage of windows with CNV signals based on the GRCh38 and T2T-CHM13 assemblies, respectively. C: Summary of CNV signals within 323 CMRGs in the T2T-CHM13 assembly. The numbers represent numbers/percentages of CMRGs, respectively. A region is likely to carry a minor allele of the global populations when CNV signals were detected at one locus in >95% of the samples. CNV signals were detected in all regions (left panel) and non-segmental duplication regions (right panel) of 323 CMRGs. D: A Vietnamese sample (HG02059) exhibited an individual-specific inversion duplication affecting the first six exons of *FLAD1* in the T2T-CHM13 assembly. Left panel: normalized depth of coverage at the *FLAD1* locus. Right Panel: alignment of one LRS read of HG02059 against the T2T-CHM13 assembly using LASTZ version 1.04.15 with default parameters (68). The gray and red lines indicate the average DoC values of protein-coding gene regions and DoC + 3*SD, respectively. E: Three African-specific duplications were identified at the *CYP4F12* locus in the T2T-CHM13 assembly. HG02011 carries a duplication (chr19:15798417-15807516) affecting the whole gene body of *CYP4F12*. HG02818 and NA19239 carry a ~1,400 bp (Chr19:15812817-15814216) duplication in intron 9 and a ~11,600 bp (chr19:15809717-15821316) duplication overlapping exons 8 to 12 of *CYP4F12*. No segmental duplication was identified in this region. The gray and red lines indicate average DoC values of protein-coding gene regions and DoC + 3*SD, respectively. F: A ~12,000 bp (Chr6:35,607,140-35,619,359) duplication affecting the whole gene body *CLPS* (not a CMRG) and the first two exons of *LHFPL5* in the T2T-CHM13 assembly was identified in samples across super-continental-populations. No segmental duplication was detected in this region. The gray and red lines indicate the average DoC values of protein-coding gene regions and DoC + 3*SD, respectively.

Given the excellent congruence of CNV analyses based on the two assemblies and a greater number of CMRGs with CNV signal detected in the T2T-CHM13 assembly compared to the GRCh38 assembly (255 versus 154, respectively), CNVs are reported based on the T2T-CHM13 assembly.

A total of 20.12% (65 out of 323), 3.10% (10 out of 323), and 46.13% (149 out of 323) of the CMRGs with CNVs were specific to individual, intra-, and inter-continental-super-populations (Figure 2C). For example, one 8,500 bp bp inversion duplication (Chr1:154,122,698-154,131,198) in a Vietnamese sample (HG02059) affected the first six exons of Flavin Adenine Dinucleotide Synthetase 1 (*FLAD1*) (Figure 2D). FAD synthase, encoded by *FLAD1*, converts the adenylation of flavin mononucleotide (FMN) to the flavin adenine dinucleotide (FAD) coenzyme and is a key enzyme in Riboflavin metabolic pathways (47). Multiple acyl-CoA dehydrogenase deficiencyies (48–50) and lipid storage myopathy due to FLAD1 deficiency have been observed to be caused by mutations in *FLAD1 (51, 52*).

Six (*ABCG8, CYP4F12, HPD, IMPA1, SSTR5*, and *TAS2R43*), one (*SLC29A4*), one (*SLC6A3*), and two (*CD247, ESRRA*) CMRGs contain CNV windows that are specific to samples from humans with African, American, South Asian, and European ancestries, respectively. For example, signals of duplication were identified at the *CYP4F12* locus in three African samples (Figure 2E). One sample carried a duplication (chr19:15798417-15807516) affecting the whole gene body of *CYP4F12*, while the other two samples carried a ~1,400 bp duplication (Chr19:15812817-15814216) in intron 9 and a ~11,600 bp duplication (Chr19:15809717-15821316) overlapping exons 8 to 12 of *CYP4F12* (transcript ID: CHM13_T0109156) (Figure 2E). *CYP4F12* (cytochrome P450 family 4 subfamily F member 12) is primarily expressed in the liver, kidney, colon, heart, and small intestine (53–55). Enzymes encoded by the members of the CYP4F family metabolize fatty acids, eicosanoids, vitamin D, and carcinogens. These enzymes are not only drug targets (56), but also catalyze the metabolism of many drugs (57–60).

A total of 149 CMRGs with CNV signals were identified in samples across continental superpopulations, suggesting that these CNV regions likely evolved in the common ancestor of modern humans or due to convergent evolution of different samples (61–63). For example, signals of duplication at the *CLPS/LHFPL5* locus were identified in the samples of humans with African, American, Asian, and European ancestries. The ~12,000 bp duplication (Chr6:35,607,140-35,619,359) affects the whole gene body of *CLPS* (that is not a CMRG) and the first two exons of *LHFPL5* (Figure 2F).

Finally, the analyses indicated that a 400 bp region (chr7:155632596-155632995) at the *DPP6* locus of the T2T-CHM13 assembly may exhibit minor alleles in the global population, as the deletion signals were identified in >95% of the samples (Supplementary Figure 2). Overall, these analyses indicate a high level of CNV polymorphisms within CMRGs.

### Short variant polymorphisms in CMRGs of global populations

In 323 CRMGs of the T2T-CHM13 assembly, a total of 152,556 short variants (119,435 SNVs and 33,121 InDels) were identified that affect 15,934,591 bp and constitute 45,964 variants per sample on average (Figure 3A). Note that some multiallelic sites contain both SNVs and InDels.

**Figure 3.**
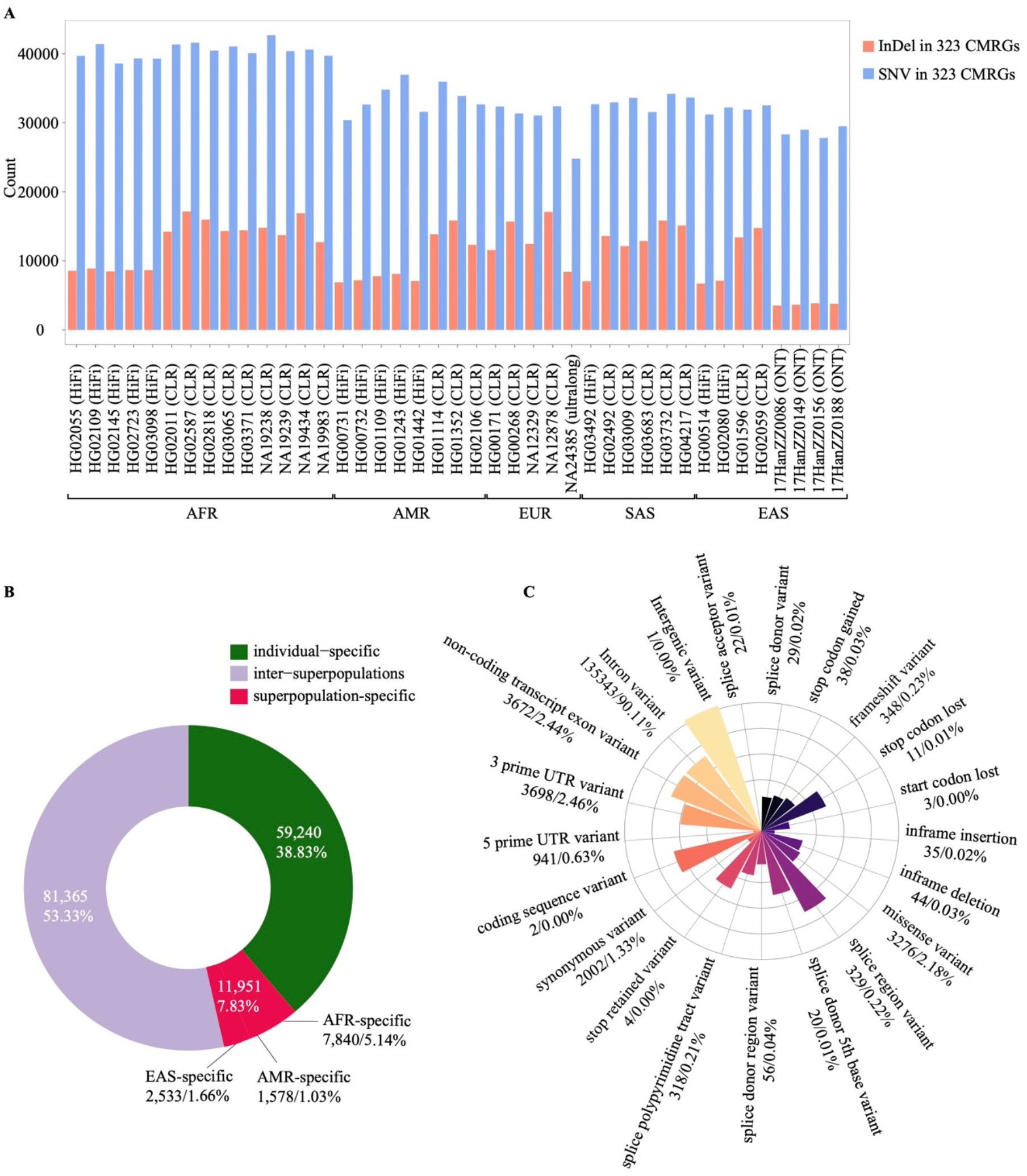
Summary of short variants detected among 323 CRMGs in the T2T-CHM13 assembly across 41 individuals using LRS data. A. Numbers of short variation (SNVs and InDels) in samples from humans with different ancestries. The X-axis indicates sample ID and sequencing technology in bracket, while the Y-axis represents numbers of SNVs and InDels. SNV: single nucleotide variant. InDel: insertion and deletion. AFR: Africans, EUR: European, AMR: American, EAS: east Asian, SAS: south Asian. B. Distribution summary of 152,556 short variants among 323 CMRGs in global populations. C. Functional annotation of 150,192 short variants in CRMGs using VEP analysis. The numbers before and after slash are the numbers and percentages of variants of each functional consequence. The most severe functional consequence of a variant was used based on the order of severity estimated by VEP when multiple consequences were predicted. The colors from light yellow to black indicate the functional consequences from low to high based on VEP.

As in the CNV analyses (Figure 2C), a large number of short variants were identified that were continental-superpopulation- or individual-specific. For example, 7,840 (5.14%), 1,578 (1.03%), and 2,533 (1.66%) short variants were specific to samples from humans with African, American, and East Asian ancestries, respectively (Figure 3B).

Functional annotation of the short variants using Ensembl VEP indicated that 95.65% of the variants were in non-coding regions such as intronic regions (90.11%), non-coding exon sequences in non-coding transcripts (2.44%), and untranslated regions (3.09%) of CMRGs (Figure 3C). Among 150,192 variants, 451 high-impact variants (based on VEP analysis) likely disrupt the normal functioning of proteins, via disruption by alternative splicing (e.g., splicing donor or acceptor sites) (51 variants in 42 genes), stop codon gain (38 variants in 30 genes), open reading frame shifts (348 variants in 156 genes), stop codon loss (11 variants in 10 genes), or start codon loss (three variants in three CMRGs).

### Side-by-side comparison of short variants in the CMRGs based on NGS and Pacbio HiFi datasets from 13 samples

Pacbio HiFi and NGS datasets were generated for 13 samples, providing an opportunity to compare the performance of short variant calling in CMRGs using both short and long reads. Short variant calling was conducted using the DeepVariant program for the NGS dataset and the PEPPER-Margin-DeepVariant program for the Pacbio HiFi dataset. An average of 5.1 million short variants per sample were detected genome-wide using the NGS data, which was ~20% less than the variants detected using the LRS data (an average of 6.5 million short variants per sample). A similar trend was also observed for CMRGs, wherein an average of 36,483 and 43,260 short variants per sample were identified based on the NGS and LRS data, respectively (Figure 4A). In particular, significantly greater numbers of short variants were identified in the segmental duplication, low complex/simply repeat regions using Pacbio HiFi data (*p* < 0.01, Wilcox two-tailed test) (Figure 4B) than when using NGS data. Variant detection is challenging in these regions using short-read data(64). In addition, 15.79% to 33.96% of the ‘high impact’ variants identified in different samples (*e.g*., splice acceptor variants, splice donor variants, and stop gains/losses) based on VEP analysis (Supplementary Table 7) could only be detected using LRS data, but not using NGS data (Figures 4C-E). For example, such variants were evident as mutations that could cause effects like a stop codon gain in *DUX4* (Figure 4C), stop codon loss in *KIR2DL3* (Figure 4D), and open reading frame shifts in *NUTM2B-AS1* (Figure 4E) and *CHMP1A* (Figure 4F).

**Figure 4.**
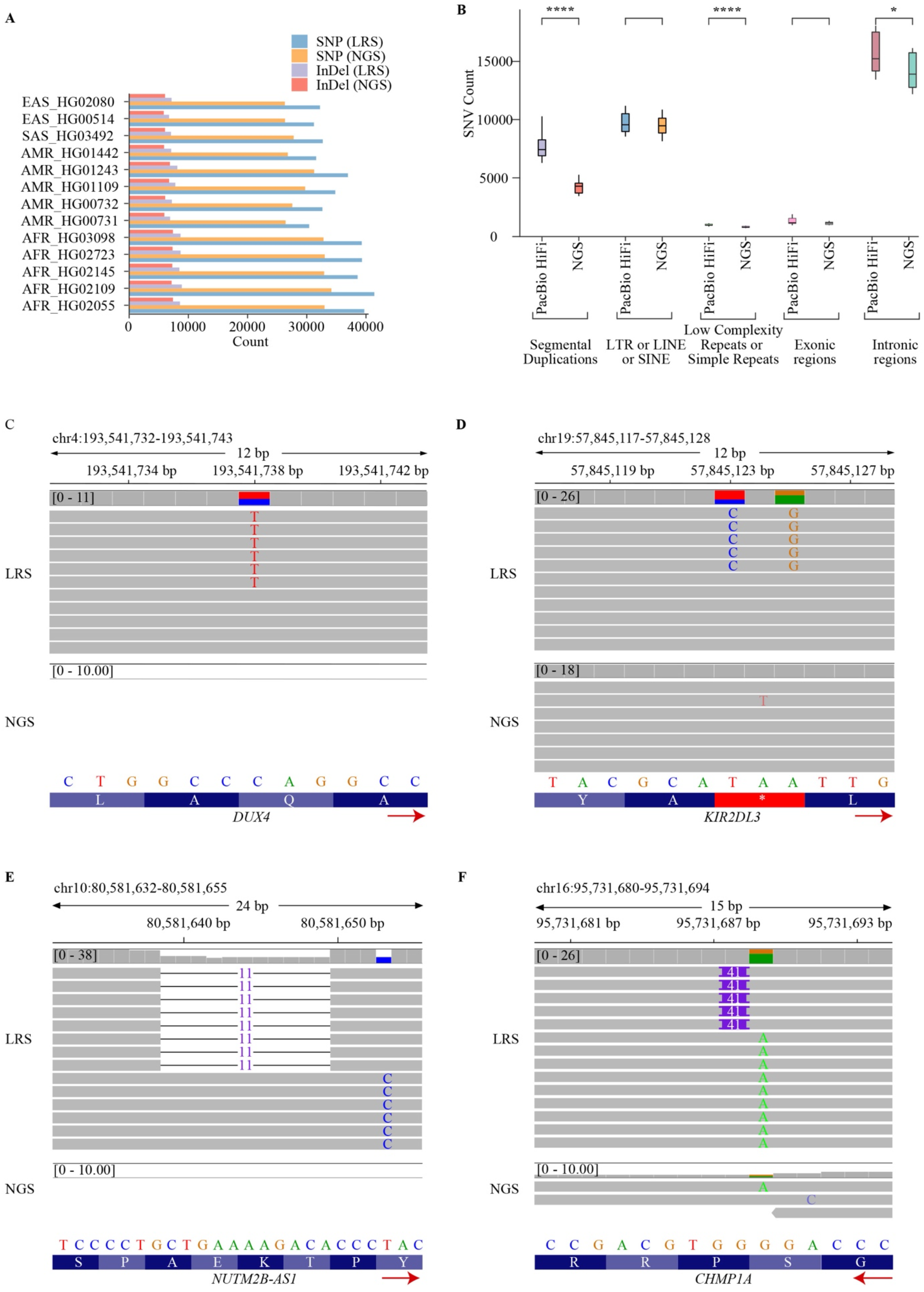
Side-by-side comparison of the short variant calls based on the NGS and LRS data of 13 individuals. A. More short variants were identified in the CMRGs per sample using Pacbio HiFi data than when using NGS data. X- and Y-axes indicate the numbers of short variants and sample ID, respectively. B. Significantly more SNVs were detected in segmental duplication, low complex/simple repeat regions using Pacbio HiFi data (*p* <0.01, Wilcox two-tailed test) than when using the NGS data. Differences were observed in the exonic, intronic, LINE, and SINE regions, but were not statistically insignificant. *: *p* <0.01, ****: *p* <0.00001 C. The C to T mutation generated a pre-stop codon in *DUX4*. This mutation was only detected using Pacbio data, since NGS reads cannot be reliably mapped to this region. D. The C to T mutation cause a stop codon loss in *KIR2DL3*. This mutation was only detected using Pacbio data. E. The GCTGAAAAGACA to G InDel generated an open reading frameshift mutation in *NUTM2B*. This InDel was only detected using PacBio data. F. A 41 bp insertion that caused an open reading frameshift mutation in *CHMP1A* was only detectable using Pacbio data. This region is difficult to identify using NGS reads, because the depth of coverage in the region considerably dropped. The red arrow indicates the direction of transcription.

## DISCUSSION

A comprehensive evaluation of CMRG genetic diversity is crucial for diagnosing genetic disorders and informing clinical treatment. However, previous studies have shown that numerous CMRGs are located in highly repetitive or complex regions of the genome, thereby hindering the detection of pathogenic variants and accurately depicting their spectrum in global populations using NGS technologies. Further, some CMRGs were misassembled in the human reference genome assemblies GRCh37 and GRCh38 (13, 25). In this study, the genetic polymorphisms of 370 CMRGs were investigated using LRS data from 41 samples collected from 19 global human populations and using the current version of the human reference genome, GRCh38 (31), along with a telomere-telomere assembly of the human genome, T2T-CHM13 (22). In addition to the potentially problematic regions in both assemblies, surprisingly large levels of genetic polymorphisms of these CRMGs were observed across human populations, indicating the need for new methods to better characterize diversity across these genes and ethnicities.

The assembly qualities of GRCh38 and T2T-CHM13 were first evaluated, with a particular focus on the 370 CMRGs. Consistent with prior studies (13, 22, 24), our analyses led to a general significant improvement in identifying protein-coding regions and CMRGs in the T2T-CHM13 assembly compared to the GRCh38 assembly, leading to a significant reduction of windows with DoC significantly deviating from the average DoC of protein-coding gene regions. Although some regions of the T2T-CHM13 genome are likely to collapse during the assembly, our analyses suggest a strong need to use T2T-CHM13 as the reference genome in future human genomic studies.

After excluding CMRGs that contain windows with abnormal DoC values in the GRCh38 and T2T-CHM13 assemblies, the diversity of SNVs, InDels, and CNVs was evaluated with 263 CMRGs and by leveraging LRS data from 41 samples of 19 globally distributed human populations. CMRGs were highly polymorphic across samples. For example, > 77% of the CMRGs in the T2T-CHM13 assembly exhibited CNV signals. Further, 188 CMRGs carried mutations that could cause severe effects including stop codon gains or losses, open reading frameshifts, or RNA splicing disruptions. Many CNVs and short variants were also observed to be individual- and super-population-specific, highlighting the necessity for considering genetic background differences among ethnic groups in future genetic testing and drug design studies. Finally, a side-by-side comparison of short variant calling using NGS and LRS data from the same individuals revealed a superior capacity of LRS data for resolving short variants in regions (*e.g*., segmental duplication and low complex/simply repeat regions) that are known to be challenging for short variant calling with NGS data. Within these regions, numerous ‘high impact’ variants were identified that could disrupt normal protein functioning. Consequently, additional LRS-based studies are needed to deepen our understanding of CMRG genetic polymorphisms.

In this study, a set of previously defined CMRGs were investigated that represent a small fraction of all medically relevant genes reported to impact various genetic disorders. Known medically relevant genes are still being discovered and thus, additional analyses are needed to characterize CMRG genetic polymorphisms using LRS data in the future. Nevertheless, some difficulties remain in using LRS data to fully resolve the genetic polymorphisms within some long repetitive regions in the T2T-CHM13 assembly (e.g., *LPA*) (44). Improvements in sequencing length, accuracy, and software algorithms for sequence data mapping and variant calling will help overcome these limitations (19, 62, 65).

An average of two samples per population was a limitation of this study leading to an inability to reliably estimate population-level site spectra. In addition, although samples from multiple ethnical groups were used, they only covered a small fraction of the genetic diversity among global populations. These factors hinder our ability to determine the prospective clinical impacts of identified variants, since rarity is used as a critical criterion for assessing the pathogenicity of a variant in clinical studies (66). In addition, many mutations were observed to be individual- and superpopulation-specific, indicating a strong need for considering differences among ethnic groups when designing drug targets. Although CNVs, SNVs, and InDels were investigated in the present study, other types of genetic alterations, like structural variants, can also interrupt normal gene functioning (67). Consequently, the results here provide a starting point for investigating the genetic diversity of CMRGs using LRS data. Advances in analytic algorithms and LRS technologies will likely bring new opportunities to achieve a deeper understanding of CMRG genetic diversity among global populations, while further improving drug development and precision medicine.

## ACKNOWLEDGEMENT

The authors thank Shuhang Li for the discussion of short variant calling.

